# Engineering circular guide RNA and CRISPR-Cas13d-encoding mRNA for the RNA editing of *Adar1* in triple-negative breast cancer immunotherapy

**DOI:** 10.1101/2025.07.22.666181

**Authors:** Shurong Zhou, Suling Yang, Jie Xu, Guizhi Zhu

**Affiliations:** Department of Pharmaceutical Sciences, College of Pharmacy, University of Michigan, Ann Arbor, MI 48109, USA; Center for Advanced Models for Translational Sciences and Therapeutics, University of Michigan Medical Center, Ann Arbor, Michigan, 48109, USA; Bioinnovations in Brain Cancer; Biointerfaces Institute; Rogel Cancer Center; Center for RNA Biomedicine. University of Michigan, Ann Arbor, MI 48109, USA

## Abstract

Clustered regularly interspaced short palindromic repeat Cas endonuclease (CRISPR-Cas) systems, such as RNA-editing CRISPR-Cas13d, are poised to advance the gene therapy of various diseases. However, their clinical development has been challenged by 1) the limited biostability of linear guide RNAs (lgRNAs) susceptible to degradation, 2) the immunogenicity of prokaryotic microorganism-derived Cas proteins in human that restrains their long-term therapeutic efficacy, and 3) off-targeting gene editing caused by the prolonged Cas expression from DNA vectors. Here, we report the development of highly stable circular gRNAs (cgRNAs) and transiently-expressing Cas13d-encoding mRNA for efficient CRISPR-Cas13d editing of target mRNA. We first optimized cgRNA for CRISPR-Cas13d editing of adenosine deaminase acting on RNA type I (*Adar1*) transcript for the combination immunotherapy of triple negative breast cancer (TNBC). cgRNAs were synthesized by enzymatic ligation of lgRNA precursors. cgRNAs enhanced biostability with comparable Cas13d-binding affinity relative to lgRNA. Next, using ionizable lipid nanoparticles (LNPs), we co-delivered the resulting *Adar1*-targeting cgRNA with an mRNA encoding RfxCas13d (mRNA-RfxCas13d), a widely used Cas13d variant, to TNBC cells. As a result, relative to lgRNA, cgRNA significantly enhanced the efficiency of *Adar1* knockdown with minimal collateral activity, which sensitized the cancer cells for cytokine-mediated cell apoptosis. In a 4T1 murine TNBC tumor model in syngeneic mice, *Adar1*-targeting cgRNA outperformed lgRNA for tumor immunotherapy in combination with immune checkpoint blockade (ICB). Collectively, these results demonstrate the great potential of cgRNA and mRNA-RfxCas13d for RNA-targeted gene editing.

## Introduction

CRISPR-Cas system has revolutionized drug development by enabling target validation, engineering cell and animal models, and gene therapy for various diseases ranging from rare genetic disorders to infectious diseases and oncology, offering a path to highly personalized and durable treatments. Among them, the CRISPR-Cas13d system is an emerging class of gene editing technologies with efficient RNA knockdown and minimal off-target effects^1^. CRISPR-Cas13d system comprises Cas13d protein as an RNA-cleaving enzyme, and a gRNA which includes a Cas13d-binding hairpin domain and a sequence complementary to the target RNA. However, the development of CRISPR-Cas13d-based RNA editing therapeutics is challenged by several limitations, including limited biostability of conventional lgRNA, inefficient delivery of protein-RNA complexes, pre-existing anti-Cas13d immunogenicity^2^ associated with the prolonged Cas13d expression, and safety concerns arising from collateral cleavage activity. lgRNA is a single-stranded RNA and has different secondary structure dependent on Cas proteins. lgRNA is prone to degradation by nucleases, resulting in limited biostability of lgRNA and consequently suboptimal gene editing efficiency and specificity. Several strategies have been tested to improve lgRNA biostability. Ribose modifications and backbone modifications are most used chemical modification to enhance nuclease resistance and allow robust in vitro and in vivo editing^3–5^. However, positional intolerance and possible toxicity might restrict its application^6,7^.

ADAR1 is a critical RNA editor that converts adenosine (A) to inosine (I) in double-stranded RNA (dsRNA). The process of A-to-I editing is essential for masking self-RNA to prevent aberrant activation of innate immune responses^8–10^, and for controlling transcriptome function by recoding mRNA^11,12^, affecting RNA stability and translation efficiency^13,14^, as well as modulating the function of microRNA^14–16^. In healthy cells, ADAR1 plays an important role in cellular viability and physiological outcomes, including proliferation, immune invasion, and differentiation^17^. ADAR1 overexpression and ADAR1 editing have been reported to be positively correlated with promoting cancer cell survival and poor cancer prognosis in several types of cancers, including oral squamous cell carcinoma^18^, triple negative breast cancer^19^, prostate cancer^20^, and non-small cell lung cancer^21^. Moreover, cancer cell ADAR1 enhances the invasiveness of cancer-associated fibroblasts^22^. Loss of ADAR1 in IFN-stimulated positive tumor cells, such as triple-negative breast cancer (TNBC)^23^, sensitizes cancer cells to cytokine-mediated cell death, primarily through interferon-stimulated gene (ISG) activation and tumor cell apoptosis^24^. These findings demonstrate ADAR1 as a potential therapeutic target for cancer immunotherapy. Indeed, ADAR1 inhibitors have been under development^20,25^, but current small-molecule-based ADAR1 inhibitors are limited by their specificity^26^.

To address these issues, here, we engineered cgRNA to enhance its biostability and used mRNA to transiently express the CRISPR-RfxCas13d protein for CRISPR-Cas13d-mediated *Adar1* knockdown in TNBC immunotherapy. Proper gRNA folding is crucial for maintaining its function, which is required for efficient Cas protein binding and target RNA recognition^27^. To enhance knockdown efficiency, we designed various gRNA sequences and identified an optimal sequence, which was subsequently circularized and codelivered with mRNA-RfxCas13d. The cgRNA demonstrated improved *Adar1* knockdown efficiency compared to its lgRNA counterpart in both mouse and human TNBC cell lines.

We also found that in mouse TNBC cells, mRNA-RfxCas13d-cgRNA mediated efficient *Adar1* knockdown, resulting in potent target cell cytotoxicity. To evaluate the therapeutic potential of this system *in vivo*, we tested the efficacy of LNP-delivered mRNA-RfxCas13d with cgRNA in a murine TNBC tumor model in syngeneic immunocompetent mice. As a result, mRNA-RfxCas13d + cgRNA, especially when combined with ICB, showed significant inhibition of tumor progression, which outperformed that of mRNA-RfxCas13d + lgRNA combined with ICB. These results demonstrated mRNA-RfxCas13d and cgRNA as a promising strategy for efficient and specific gene editing, with implications for application in other gene editing systems.

## Results

### *Adar1* gRNA structural optimization and cgRNA synthesis

The gRNA for RfxCas13d comprises a direct repeat (DR) region and an RNA-targeting spacer sequence, which bind to RfxCas13d protein and its RNA target, respectively. The optimal length of spacer sequence is reported to be 23-30 nucleotides (nts) for efficient target RNA knockdown, and changes in the DR region could completely abrogate target knockdown^27^. Since gRNA folding is critical, we optimized the sequence by adding extra nts at the 3’ end of the spacer sequence to elongate the overall gRNA length for circularization while maintaining its original secondary structure. We tested three different gRNA sequences targeting *Adar1* in 4T1 murine TNBC cells: the original gRNA sequence, a precursor gRNA that featured two direct repeats (DR) on both ends, and a gRNA with an extra 10 non-target nts. 24 h after transfection of 4T1 cells with gRNA and mRNA-RfxCas13d using Lipofectamine transfection reagent, the *Adar1* RNA transcript levels were measured by RT-qPCR. The gRNA with extra 10 non-target nts showed the most efficient *Adar1* knockdown efficiency (Figure 1A). Worth noting, the predicted minimum free energy (MFE) of the corresponding gRNA (Figure S1) showed no correlation with the knockdown efficiency reported previously using plasmid delivery system.^27^ Such discrepancy is presumably because our CRISPR-RfxCas13d system based cgRNA and mRNA-RfxCas13d may be impacted by differential factors compared to the previous systems.

**Figure 1.**
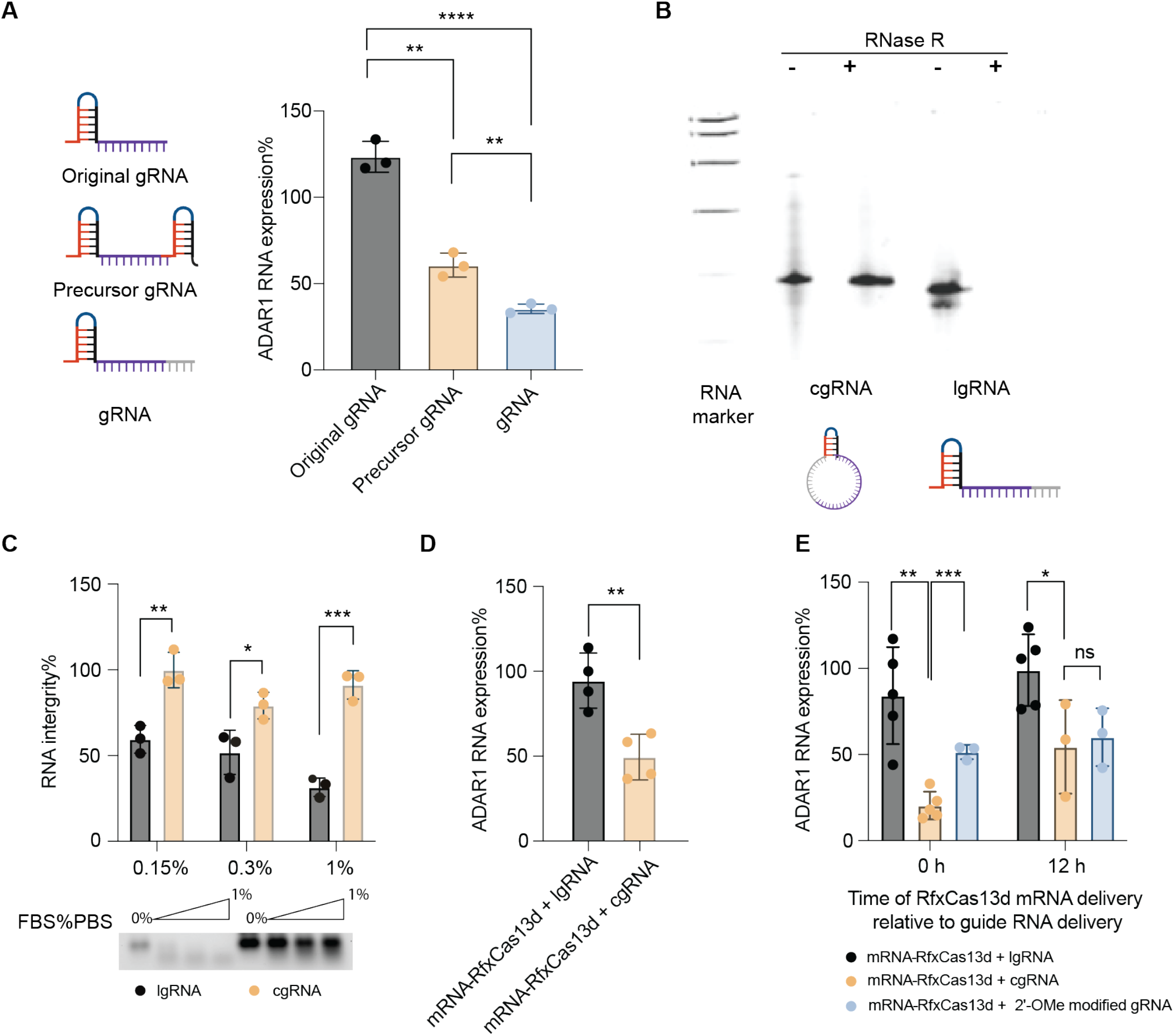
Relative to lgRNA, cgRNA showed enhanced biostability and improved *Adar1* knockdown efficiency in cells. A) Left: predicted secondary structure of three different engineered gRNA; right: *Adar1* RNA levels of 4T1 cells 24 h after transfection with mRNA-RfxCas13d (1 μg/mL) with different gRNA (1 μg/mL). RNA was transfected using Lipofectamine 2000. B) PAGE gel electrophoresis of lgRNA and cgRNA before and after RNase R treatment (15 min). C) RNA integrity analysis of lgRNA and cgRNA based on agarose gel of RNA treated with diluted FBS. Bottom: a representative agarose gel image; top: summarized fractions of intact RNA based on three independent replicates of agarose gel results. D) *Adar1* RNA levels in 4T1 cells 24 h after treatment with mRNA-RfxCas13d (0.5 μg/mL) and lgRNA or cgRNA (0.25 μg/mL) at low concentrations. E) *Adar1* RNA levels in 4T1 cells 24 h after treatment with mRNA-RfxCas13d with lgRNA, cgRNA or 2’-OMe modified gRNA. Data: mean ± SD. n.s.: not significant, p> 0.05; *p < 0.05, **p < 0.01; ***p < 0.001. p value was determined by an unpaired student t-test.

After determining the gRNA sequence, we performed gRNA circularization using T4 RNA ligase to enhance gRNA stability. In denaturing polyacrylamide gel electrophoresis (PAGE), the resulting cgRNA showed a slightly slower migration rate compared to the corresponding lgRNA with the identical sequence as cgRNA, which verified the successful gRNA circularization; moreover, relative to lgRNA, cgRNA demonstrated enhanced resistance to exonuclease RNase R treatment (Figure 1B). Successful circularization was further verified by Sanger sequencing of the complementary DNA (cDNA) for the ligation junction. Specifically, cgRNA, but not lgRNA, yielded amplified cDNA fragments following reverse transcription and PCR (Figure S2), and Sanger sequencing of the resulting cDNA verified accurate ligation at the circularization site. Due to the absence of free 5’ and 3’ termini, cgRNA is expected to be more stable than its linear counterpart. To validate this, we compared the stability of cgRNA and lgRNA in PBS containing a series of concentrations of fetal bovine serum (FBS) at 37 °C for 15 minutes. RNA integrity analysis by gel electrophoresis revealed over 90% intact cgRNA under these conditions, in contrast to approximately 50% intact lgRNA, suggesting the superior ability of cgRNA to resist exonuclease degradation and its resulting superior biostability (Figure 1C).

### Using mRNA-RfxCas13d, cgRNA enhanced *Adar1* knockdown efficiency relative to lgRNA in cells

We then evaluated the *Adar1* knockdown efficiency of cgRNA and lgRNA in 4T1 cells. To test the gRNA dose-dependent effect, we used two different gRNA concentrations with a constant concentration of mRNA-RfxCas13d. At gRNA concentration of 0.5 μg/mL, both lgRNA and cgRNA achieved comparable knockdown efficiency (Figure S3). At the gRNA concentration of 0.25 μg/mL, the knockdown efficiency of cgRNA reached ca. 45%, which is 2.2-fold of that for lgRNA (Figure 1D). This suggests that lgRNA is susceptible to degradation, resulting in insufficient levels to bind to RfxCas13d protein and mediate target gene knockdown. In contrast, the stability of cgRNA likely contributes to its sustained activity and prolonged knockdown effects. Additionally, we compared the performance of lgRNA, cgRNA, and a benchmark gRNA with 2’-O-methyl (2’-OMe) modification commonly used to enhance gRNA stability and editing efficiency for CRISPR-Cas systems^28,29^. When co-delivered with mRNA-RfxCas13d, cgRNAs exhibited the highest knockdown efficiency among all these gRNAs (Figure 1E). Moreover, in sequential transfection, where gRNA was introduced first and mRNA was added 12 h later, cgRNA showed comparable knockdown efficiency to that of the 2’-Ome-modified gRNA. This suggests that cgRNA enhances the stability of linear gRNA and hence improves its target gene knockdown efficiency.

### With an intact circular structure, cgRNA was avidly bound with RfxCas13d protein

To investigate the potential impact of gRNA circularization on the gRNA binding with RfxCas13d protein, we used a recombinant RgxCas13d protein to assemble gRNA-RfxCas13d ribonucleoprotein complexes for cgRNA or lgRNA, respectively. PAGE-based electrophoretic mobility shift assay (EMSA) revealed that lgRNA and cgRNA exhibited comparable binding ability to RfxCas13d protein (Figure 2A), indicating that RNA circularization did not adversely affect cgRNA’s binding ability with RfxCas13d protein. This provides the basis for cgRNA to mediate target RNA-specific CRISPR-Cas13d editing.

**Figure 2.**
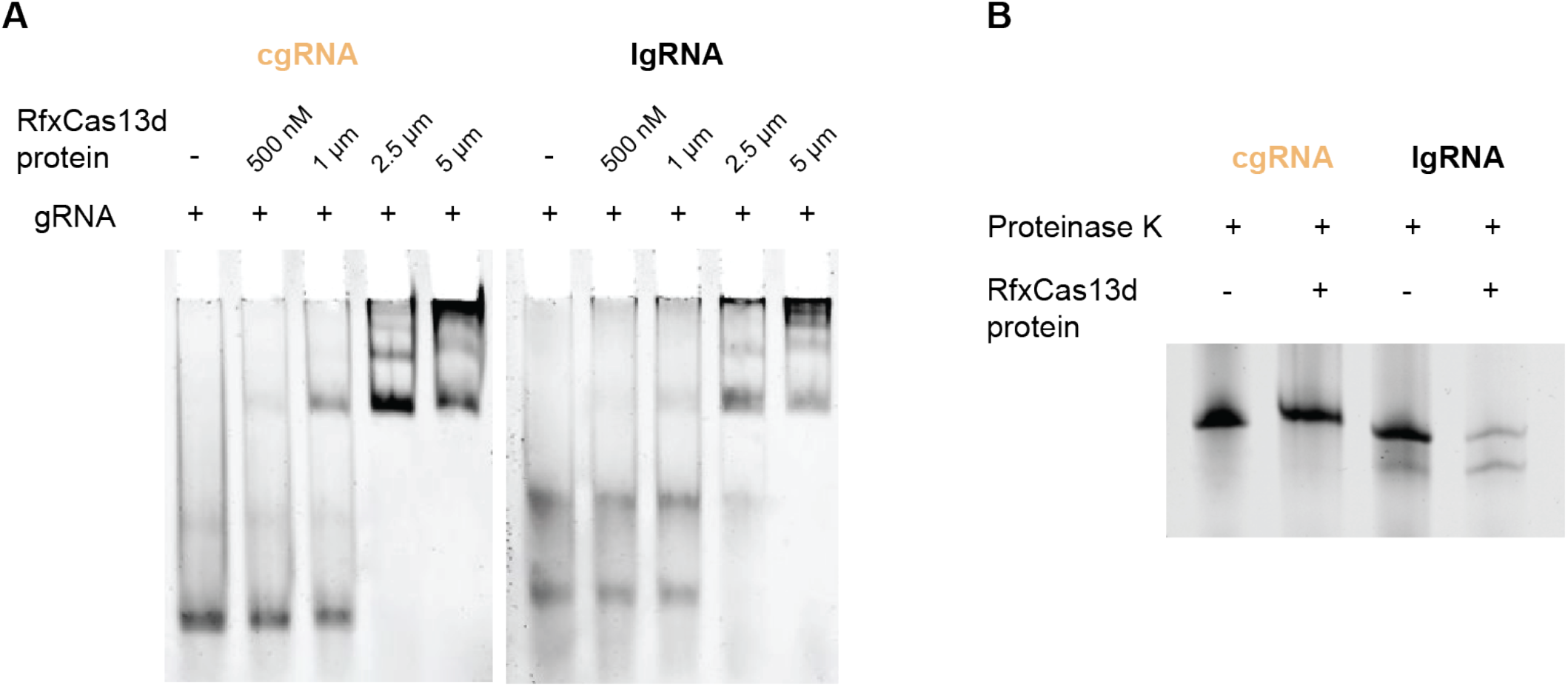
cgRNA showed avid binding with RfxCas13d protein while maintaining the intact circular RNA structure. A) EMSA results of RfxCas13d protein mixed with cgRNA and lgRNA, respectively. RNA concentration: 1 μM. B) Gel electrophoresis image showing the results of cgRNA and lgRNA co-incubated with RfxCas13d protein for 45 min. This RNA cleavage assay suggested that cgRNA maintained its circular RNA structure when bound with RfxCas13d protein, without being processed by RfxCas13d protein into linear RNA.

Given the intrinsic self-cleavage ability of the Cas13d endonuclease to process its own gRNA,^30^ we further evaluated the structural integrity of the cgRNA that was complexed with recombinant RfxCas13d protein *in vitro*. The RNA cleavage assay showed that the cgRNA remained intact, with undetectable processing into linear RNA after incubation with RfxCas13d protein for 45 min (Figure 2B). These findings suggest that, when bound with RfxCas13d protein, cgRNA still maintains its circular structure and likely functions in its circularized form during target RNA editing, which is expected to avoid early linearization of cgRNA and maximize the biostability of cgRNA.

### cgRNA + mRNA-RfxCas13d mediated transient and specific *Adar1* knockdown to sensitize TNBC cells for cytokine-mediated cell apoptosis

We further investigated the downstream effects of *Adar1* knockdown in 4T1 cells. It has been reported that ADAR1 depletion in ADAR1-overexpressing cancer cells, including TNBC cells, can inhibit cancer cell proliferation and increase tumor cell sensitivity to immunotherapy. In our study, a reduction in ADAR1 protein was observed in 4T1 cells 2 days post co-transfection of cgRNA + mRNA-RfxCas13d, reaching as low as 37% around 3 days post transfection, and then returning to baseline by 5 days post transfection (Figure 3A). A similar ADAR1 protein reduction level was observed in MC38 murine colorectal cells that were treated with cgRNA + mRNA-RfxCas13d as above (Figure S4). We then studied the impact of *Adar1* knockdown by cgRNA + mRNA-RfxCas13d on the sensitivity of cytokine-mediated cell apoptosis, which is critical for cancer immunotherapy. Two days after transfection of 4T1 cells with cgRNA + mRNA-RfxCas13d, the resulting cells were additionally treated with interferon gamma (IFN-γ) for another 3 days. IFN-γ is a cytokine that plays a crucial role in tumor cell killing for many types of cancer immunotherapy, including ICB and adoptive T cell therapy. Compared to controls, treatment of mRNA-RfxCas13d and cgRNA promoted the apoptosis and reduced the cell viability of 4T1 cells (Figure 3B-C). To ensure that the observed cell toxicity was not due to collateral RNA cleavage by the CRISPR-Cas13d system, we assessed 4T1 cell viability in the absence of IFN-γ treatment for 48 hours. No reduction in cell viability was observed in cells transfected with mRNA-RfxCas13d alone or together with gRNA (Figure S5), indicating minimal nonspecific cell toxicity associated with collateral RNA cleavage. To further investigate the possibility of collateral activity by mRNA-RfxCas13d and cgRNA, we then used human TNBC MDA-MB 231 reporter cells expressing both GFP and firefly luciferase. Treatment of these cells with *Adar1*-targeting cgRNA together with mRNA-RfxCas13d did not alter GFP (Figure 3D) or luciferase expression (Figure S6). This further verified the undetectable Cas13d-associated collateral RNA cleavage activity. According to the DepMap database^31^, *Adar1* is expressed at a medium-copy number in cancer cell lines (Figure S7), which may account for the lack of observable collateral activity in this context. Overall, these results demonstrated the ability of cgRNA + mRNA-RfxCas13d to mediate transient and specific *Adar1* knockdown in TNBC cells, resulting in the immunosensitization of these cells for apoptosis.

**Figure 3.**
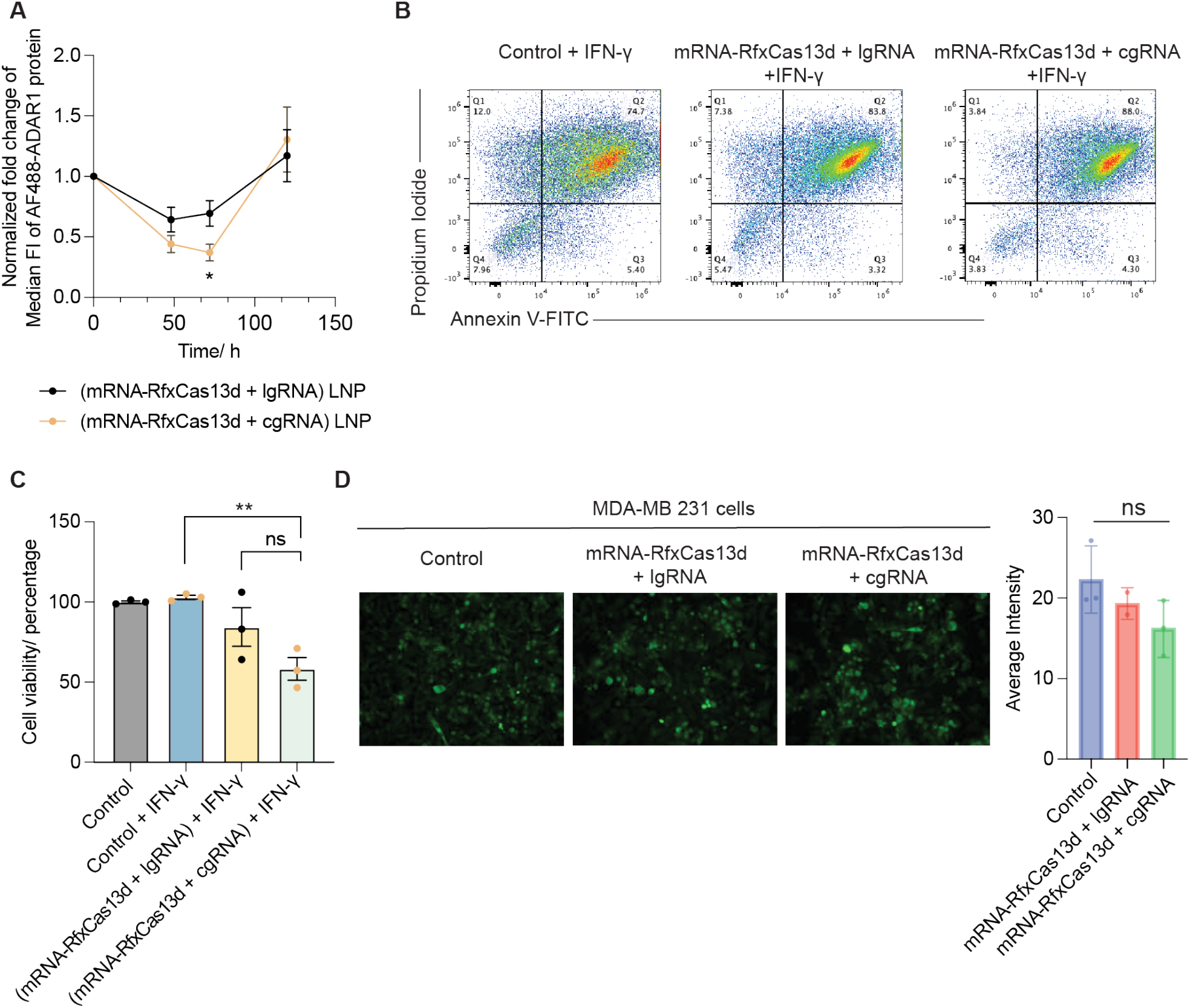
cgRNA + mRNA-RfxCas13d mediated transient and specific *Adar1* knockdown with minimal collateral activity, leading to increased TNBC cell apoptosis. A) ADAR1 protein expression level in 4T1 cells analyzed by flow cytometry. B) Annexin V staining of IFN-γ-treated 4T1 cells after *Adar1* knockdown. C) Cell viability test after *Adar1* knockdown following IFN-γ treatment in 4T1 cells. D) Evaluation of RfxCas13d-mediated EGFP expression by collateral activity in human TNBC cells. Data: mean ± SD. n.s.: not significant, p> 0.05; *p < 0.05, **p < 0.01. p value was determined by an unpaired student t-test.

### LNP co-delivery of mRNA-RfxCas13d and cgRNA sensitized 4T1 tumor to ICB therapy in mice

ADAR1 is an endogenous RNA base editor that plays a critical role in RNA editing, which is essential to cell immune homeostasis by preventing abundant cytosolic RNA from eliciting overly strong proinflammatory innate immune responses in normal cells^33^.To evaluate the therapeutic efficacy of the *Adar1* knockdown effect mediated by mRNA-RfxCas13d and cgRNA, we codelivered mRNA-RfxCas13d and *Adar1* cgRNA using SM-102 LNPs^32^. We first evaluated the biodistribution of LNP encapsulation a reporter mRNA encoding enhanced luciferase protein (mRNA-fLuc) after a single-dose intratumoral treatment in 4T1 tumor-bearing syngeneic Balb/c mice. IVIS imaging of the resulting mice showed LNPs encapsulating mRNA-fLuc was specifically expressed in tumor tissue for at least 3 days (Figure 4A). Further, we used LNPs encapsulating another mRNA encoding EGFP (mRNA-EGFP) to study the intratumoral cellular distribution of mRNA expression upon tumor treatment as above. As shown by flow cytometric analysis of EGFP signal in tumor-derived single cells, mRNA-EGFP was primarily expressed in CD45^-^ cells, especially in CD45^-^ EpCAM^+^ cells, which are likely 4T1 tumor cells^34,35^ (Figure 4B).

**Figure 4.**
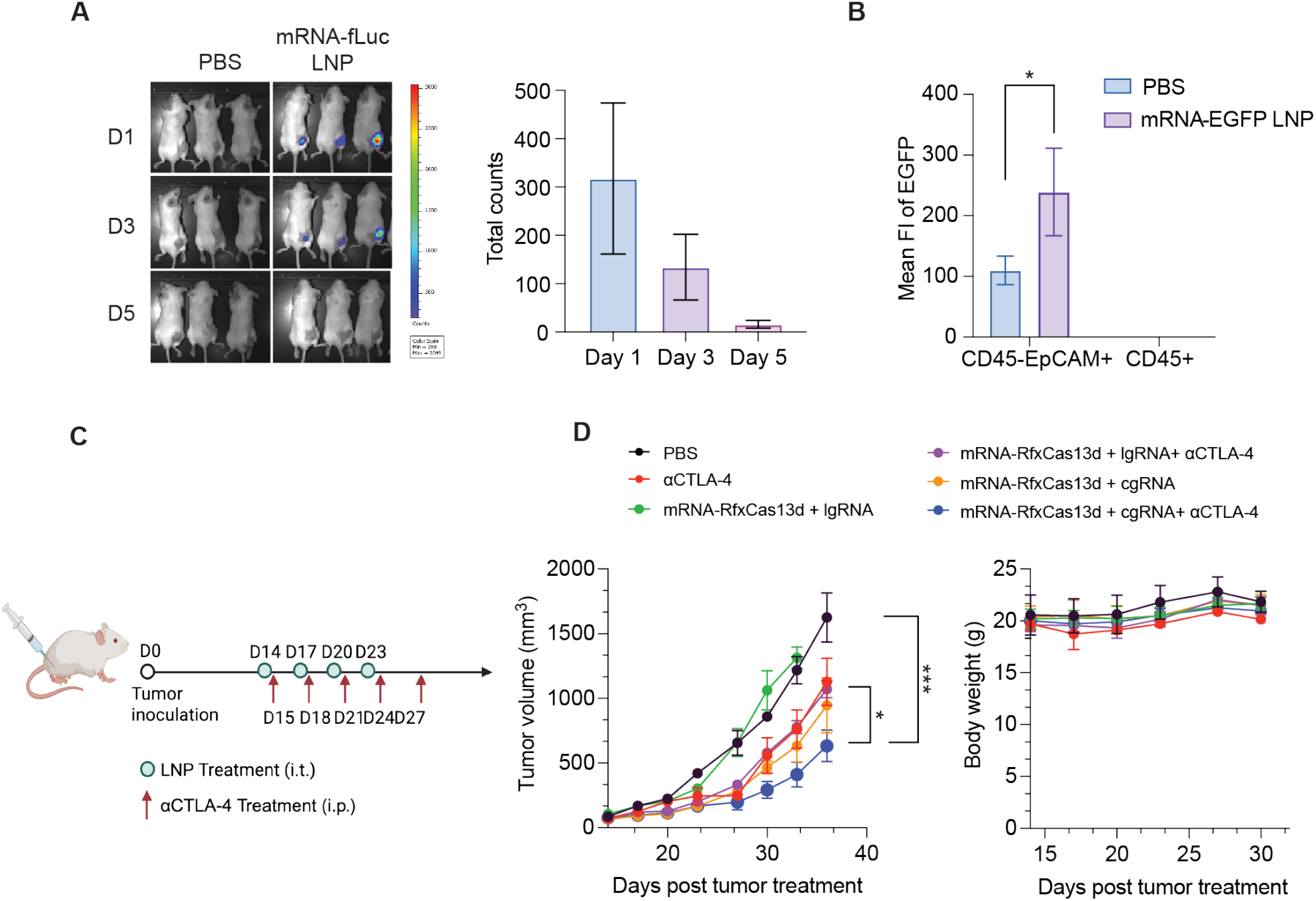
LNP co-delivery of mRNA-RfxCas13d and cgRNA for ICB combination immunotherapy of 4T1 tumors in mice. A) IVIS imaging of mRNA-fLuc expression in 4T1 tumors after a single-dose intratumoral injection of mRNA-fLuc LNPs (5 μg RNA /mouse). B) Flow cytometric analysis of EGFP expression in different cell populations in 4T1 tumors delivered by mRNA-EGFP LNPs (5 μg RNA /mouse). C) Timeline of 4T1 tumor therapy study. D) Tumor growth curves and mouse body weights after the indicated treatments. Data: mean ± SEM. n.s.: not significant, p> 0.05; *p < 0.05, **p < 0.01; ***p < 0.001. p value was determined by one-way ANOVA.

We further investigated the tumor therapeutic efficacy of the above LNPs co-loaded with mRNA-RfxCas13d and *Adar1* cgRNA, alone and in combination with ICB, in the above 4T1 tumor model. Again, 4T1 tumor cells were subcutaneously injected at the flank of Balb/c mice. When the tumor sizes reached approximately 70 mm^3^, the mice were intratumorally treated with the above RNA-loaded LNPs and controls for four times at 3-day intervals, accompanied by five doses of intraperitoneal anti-cytotoxic T-lymphocyte-associated protein 4 (αCTLA-4) treatment (Figure 4C). Relative to αCTLA-4 alone, mRNA-RfxCas13d and cgRNA markedly promoted the tumor inhibition efficacy by αCTLA-4; moreover, cgRNA outperformed lgRNA to sensitize tumors for ICB therapy (Figure 4D). No mouse body weight loss was observed during any of these treatments, which suggests the promising safety of LNP-delivered mRNA-RfxCas13d and cgRNA. Overall, these results demonstrated the benefit of gRNA circularization for *Adar1* knockdown to sensitize TNBC cells for ICB immunotherapy.

## Discussion

The development of CRISPR-Cas systems has significantly expanded the toolkit for gene editing and gene therapy. Yet, the clinical application of many CRISPR-Cas systems has faced multiple challenges. First, lgRNA often have limited biostability due to their susceptibility to enzymatic degradation and hydrolysis, resulting in suboptimal gene editing efficiency. Chemical modification for gRNA, including backbone modification using phosphorothioate and phosphonoacetate, as well as ribose modification using 2’-Ome and 2’ fluoro, have been studied to increase nuclease resistance and gene editing efficiency. However, these modifications are not only often costly but may also impair the binding affinity of gRNA with Cas protein and increase the frequency of off-target gene editing. Second, in vertebrates, prolonged expression of bacteria-derived Cas proteins may elicit anti-Cas immunity and cause off-targeting effects, both of which may restrain their long-term therapeutic efficacy. Lastly, viral vectors (e.g., adeno-associated viruses) widely used for in vivo delivery of CRISPR-Cas gene editing systems are limited by their dose-dependent immunogenicity and the potential genotoxicity associated with unintended genomic integration.

To address these challenges, we developed highly stable cgRNAs and transiently expressing mRNA-RfxCas13d, which are co-delivered by LNPs for efficient CRISPR-Cas13d editing of target *Adar1* transcript in TNBC immunotherapy. Synthesized by a one-step intramolecular enzymatic ligation, cgRNAs showed enhanced biostability and comparable Cas13d-binding affinity relative to lgRNA. As a result, relative to lgRNA, cgRNA promoted *Adar1* knockdown efficiency in murine and human TNBC cells with minimal collateral activity, which sensitized the cancer cells for cytokine-mediated cell apoptosis. Emerging non-viral delivery systems, such as LNPs that can deliver protein-RNA complexes or co-deliver Cas-encoding RNA and gRNA, offer a promising alternative by enabling transient Cas exposure and potentially improving safety profiles. We then used LNPs to co-deliver *Adar1*-targeting cgRNA mRNA-RfxCas13d to tumor cells upon intratumoral administration in a TNBC murine tumor model in syngeneic mice. cgRNA outperformed lgRNA for ICB combination tumor immunotherapy. These results demonstrate the potential of LNPs of cgRNA + mRNA-RfxCas13d for efficient nonviral gene editing in the treatment of various diseases.

This study also has several limitations. First, though we tested collateral activity of using mRNA-RfxCas13 and cgRNA in vitro, the safety profile of LNP-delivered mRNA-RfxCas13 and cgRNA, including the immunogenicity against RfxCas13 and toxicity, remains to be comprehensively evaluated. Moreover, while we demonstrated the therapeutic efficacy of this approach by intratumoral administration of (cgRNA + mRNA-RfxCas13d) LNPs, this administration route can be limited to application in surgically accessible tumors. Therefore, alternative administration routes, such as intravenous administration, warrant testing to broaden the application of these therapeutics. Lastly, it remains to be evaluated whether gRNA circularization can be effectively applied to other CRISPR systems, such as CRISPR-Cas9 or prime editors, which utilize more complex gRNA structures. Further efforts in rational gRNA sequence design, along with the purification and characterization of circularized gRNAs, will be crucial to broaden the applicability of this strategy across various CRISPR-based technologies.

## Materials and Methods

### Cell culture

4T1 cells and MDA-MB 231 luciferase-EGFP dual reporter cells, were cultured in RPMI 1640 medium. MC38 cells were cultured in DMEM medium. Medium was supplemented with 10% FBS and 0.1% penicillin and streptomycin. All cells were cultured in a Biosafety Level II incubator (5% CO_2_, 37 °C).

### In vitro transcription of mRNA-RfxCas13d

mRNA-RfxCas13d was synthesized by in vitro transcription using T3 polymerase. The linearized plasmid (Addgene: 141320) was used as a DNA template and in vitro transcribed using MEGAscript™ T3 Transcript kit (Invitrogen™), following the manufacturer’s protocol. UTP was substituted with pseudo-UTP and Cap Analog (m7G(5’)ppp(5’)(2’-oMeA)pG) was incorporated as a cap analog. The reaction was incubated at 37 °C for 2 h, followed by adding 1 μL Turbo DNase to remove the DNA template. The synthesized mRNA was purified using RNA clean-up column (Zymo Research), and the mRNA integrity was confirmed using gel electrophoresis.

### gRNA circularization

Full-length linear gRNA precursors were synthesized by Hippo Bio. gRNA circularization was achieved using T4 RNA ligase I. In brief, gRNA was diluted in T4 RNA ligase I reaction buffer and heated at 90°C for 5 min,and was then cooled down to 25 °C gradually. T4 RNA ligase I (NEB), 10 mM ATP, and 50% PEG 8000 was added following the guidance from the manufacturer. The circularized gRNA was purified using an RNA clean-up column (Zymo Research) and concentrated by RNase R (Biosearch Technologies) and verified by 10% denatured PAGE gel electrophoresis.

### In vitro assessment of *Adar1* knockdown

To evaluate *Adar1* RNA levels, 4T1 cells were seeded in a 12-well plate at a cell density of 3 × 10^5^ cells / well. mRNA-RfxCas13d and *Adar1*-targeting cgRNA or lgRNA were transfected using Lipofectamine 2000. Total RNA from the cells was extracted 24 hours post-transfection for RT-qPCR using the SYBR green assay (Applied Biosystem). For ADAR1 protein expression evaluation, the transfected cells were collected at different time points for ADAR1 immunostaining using Alexa Fluor 488-labeled anti-ADAR1 antibody (Santa Cruz), followed by flow cytometric analysis of cell fluorescence intensities.

### RNA stability

Equal amounts of lgRNA and cgRNA were dispensed in PBS with different fraction of FBS (0, 0.15%, 0.3%, 1% v/v). After 15 min of incubation at 37°C, equal volumes of each sample were collected for RNA integrity analysis using 2% TBE agarose gel electrophoresis. The integrity of each gRNA sample was quantified by Image Lab (Bio-Rad) and normalized to the corresponding RNA sample incubated in PBS without FBS (0%).

### RT-PCR of cgRNA and cDNA Sanger sequencing

RT-PCR primers were designed to amplify the ligation junction site of cgRNA. RT-PCR was performed using OneStep RT-PCR kit (Qiagen), with either lgRNA or cgRNA as the RNA template. The resulting PCR products were analyzed by 2% TAE agarose gel electrophoresis. The smear band observed in the circular RNA was excised from the gel, purified by a DNA gel recovery kit (Zymo Research), and subject to Sanger Sequencing.

### EMSA

RfxCas13d protein was expressed and purified by the Center for Structure Biology at the University of Michigan. The purified RfxCas13d protein was mixed with gRNA in the binding buffer (2 mM MgCl_2_, 50 mM Tris) at different molar ratios on ice. The resulting mixture was incubated at room temperature for 30 min. The formation of the RfxCas13d protein-gRNA complex was assess using a 6% native PAGE gel.

### RNA cleavage assay

RfxCas13d protein and gRNA were mixed at a 2:1 molar ratio in RNA Cleavage Buffer (25 mM Tris pH 7.0, 1 mM DTT, 5 mM MgCl_2_). The reaction was prepared on ice and incubated at 37 °C for 45 min. The reaction was subsequently incubated at 37 °C for 45 min and quenched with 10 μL enzyme stop solution (10 mg/mL Proteinase K, 4 M Urea, 80 mM EDTA, 20 mM Tris pH 8.0) at 37 °C for 15 min. Then 2x RNA loading buffer was added for RNA denaturation at 90 °C for 5 min. RNA samples were run in 12% 7.5 M Urea PAGE gel at 220 V for 40 min.

### Collateral activity

MDA-DB 231 cells expressing both luciferase and EGFP were seeded in a 96-well plate at a cell density of 1 × 10^4^ cells / well. Cells were transfected with mRNA-RfxCas13d and *Adar1*-targeting gRNA using Lipofectamine 2000. EGFP expression in cells was assessed under a fluorescence microscope, and luciferase expression was measured using One-Glo™ luciferase assay kit (Promega), respectively, 48 hours post transfection.

### Cell death assay

4T1 cells were seeded in 12-well plates at a cell density of 3 × 10^5^ cells / well. mRNA-RfxCas13d and *Adar1*-targeting cgRNA or lgRNA were transfected using Lipofectamine 2000. IFN-γ (200 ng/mL) was added 48 hours post transfection and incubated for an additional 72 hours. Cell viability was assessed using the CellTiter-Glo™ luciferase assay (Promega), and apoptosis was evaluated by flow cytometry following staining with the FITC-Annexin V Apoptosis Detection Kit (BioLegend).

### Synthesis of RNA-loaded LNPs

mRNA-RfxCas13d and gRNA were formulated in LNPs using the ethanol dilution method. In brief, SM-102, cholesterol, DSPC, and DMG-PEG2000 were dissolved in ethanol and RNA was dissolved in 10 mM sodium citrate buffer. All lipids were mixed with a molar ratio of 50: 38.5: 10: 1.5. LNPs were formulated via NanoGenerator Flex-S (PreciGenome), at an aqueous to ethanol ratio of 3: 1(v/v) with a final mass ratio of 20: 1. The fresh LNPs were dialyzed using a Pur-A-Lyzer™ dialysis kit (Sigma-Aldrich, MWCO: 6 KDa) at 4 °C overnight, and concentrated by ultracentrifugation tubes (Millipore, MWCO: 100 K) for in vivo injection. LNPs were diluted in 1× DPBS and measured using Zetasizer (Malvern Panalytical) to determine sizes and zeta potential.

### Animal study

Balb/c mice (6–8 weeks old) were purchased from Charles River Laboratory. All animals were maintained at the University of Michigan’s animal facilities under specific pathogen-free conditions and treated in accordance with the regulations and guidelines of the Institutional Animal Care and Use Committee. The University of Michigan Institutional Animal Care and Use Committee approved all animal experiments.

### Biodistribution and mRNA expression in mice by IVIS imaging

To study tissue biodistribution for mRNA-fLuc LNPs, mRNA-fLuc LNPs were injected intratumorally into 4T1 tumor-bearing Balb/c mice. After 6 h, the mice were intraperitoneally injected with D-luciferin (10 mg/kg) (Goldbio), followed by bioluminescence imaging of the mice 15 minutes later. To study cellular uptake for mRNA-EGFP LNPs, mRNA-EGFP LNPs were injected intratumorally into 4T1 tumor-bearing Balb/c mice. After 6 h, the mice were sacrificed, and the injected tumor was dissected for flow cytometric analysis of EGFP expression in different cell populations.

### Tumor therapy

Tumors were established by subcutaneous injection of 3 × 10^5^ tumor cells into the right flank of mice. Treatment was initiated when the average tumor size reached 100 mm^3^. LNPs co-loaded with mRNA-RfxCas13d and gRNA were intratumorally injected at a total dose of 15 μg RNA (mRNA/gRNA = 1:2 in mass ratio) into tumors every 3 days for 4 times. For ICB, mice were intraperitoneally injected with αCTLA-4 at a dose of 100 μg every 3 days for 5 times. Tumor sizes and mouse body weight, as well as health conditions, were monitored. The humane endpoint for mice was defined as either a 20% drop in mouse body weight or a tumor size of 1500 mm^3^.

## Acknowledgements

G.Z. acknowledges funding support from NIH (R01CA286122, R01CA266981, R01AI168684, R35GM143014), DoD CDMRP Breast Cancer Breakthrough Award Level II (BC210931/P1), American Cancer Society Research Scholar Grant (RSG-22-055-01-IBCD), University of Michigan Rogel Cancer Center Discovery Award and Forbes Institute for Cancer Discovery Award. Research reported in this publication was supported by the National Cancer Institute under Award Number P30 CA046592 by the use of the following Cancer Center Shared Resource: Flow Cytometry, Immune Monitoring, Pharmacokinetics, and Proteomics. We thank the University of Michigan ULAM (Unit for Laboratory Animal Medicine) In Vivo Animal Core (IVAC) and the Center for Structural Biology for providing service. The content is solely the responsibility of the authors and does not necessarily represent the official views of the National Institutes of Health.

## Author contributions

S.Z., G.Z. conceptualized the project. All authors reviewed the manuscript.

