## Supplementary Information for "Engineering circular guide RNA and CRISPR-Cas13d-encoding mRNA for the RNA editing of *Adar1* in triple-negative breast cancer immunotherapy"

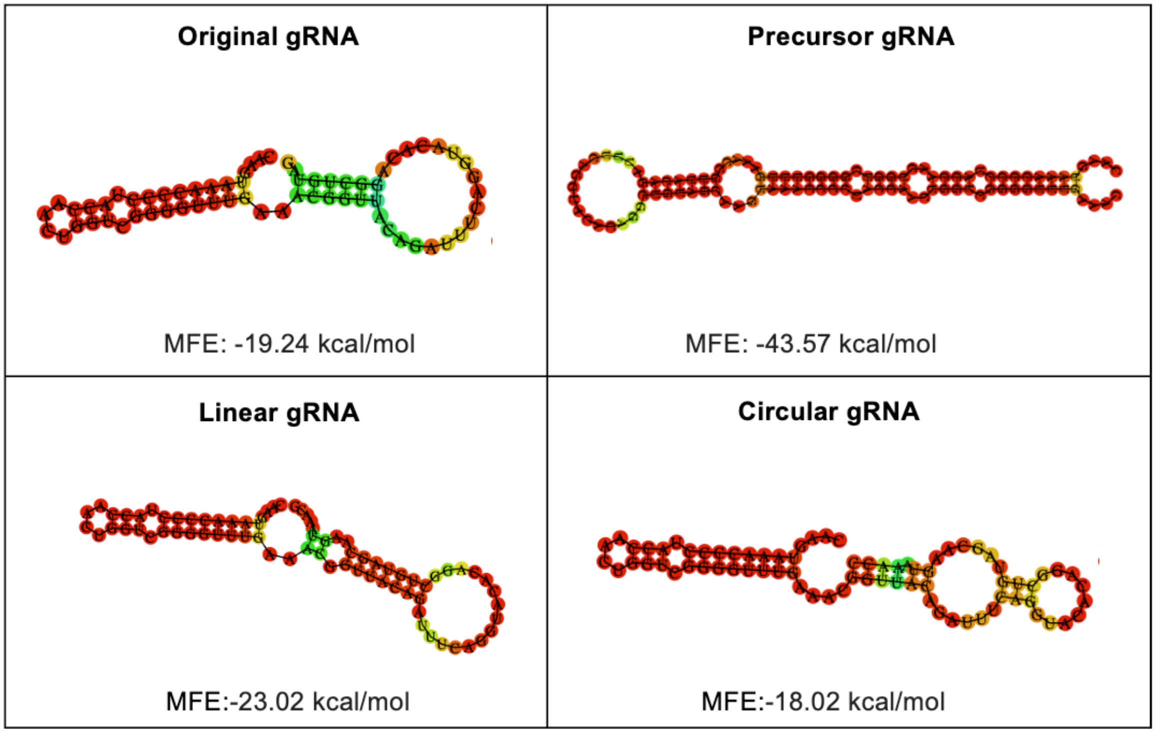


**Figure S1.** Predicted secondary structures of RNA sequences using RNAfold (http://rna.tbi.univie.ac.at/cgi-bin/RNAWebSuite/RNAfold.cgi).

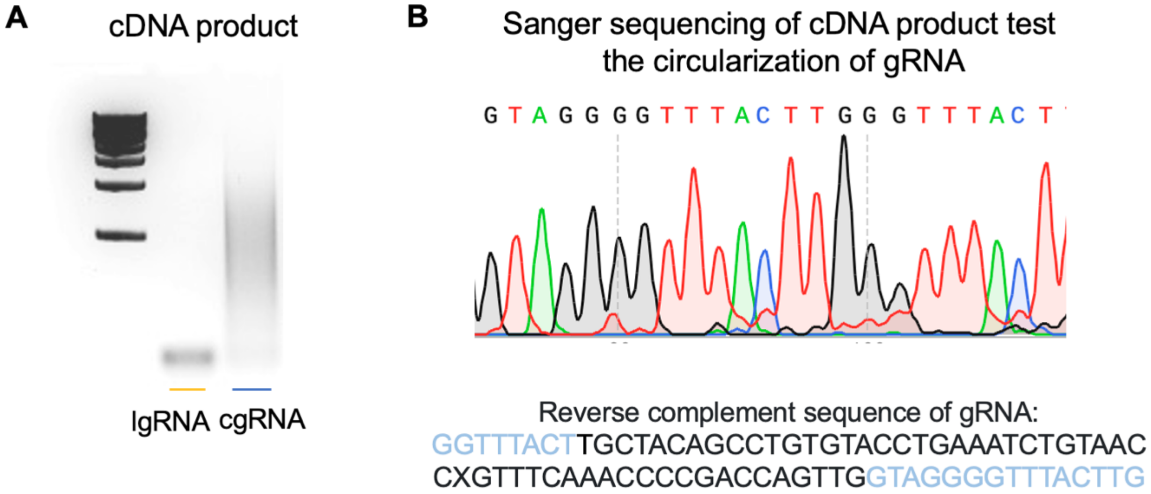


**Figure S2**. Sanger sequencing of the cDNA of cgRNA junction site validates cgRNA synthesis. (A) An image of agarose gel electrophoresis for reverse-transcribed cDNA of cgRNA junction site. (B) Sanger sequencing results of the above reverse-transcribed cDNA.

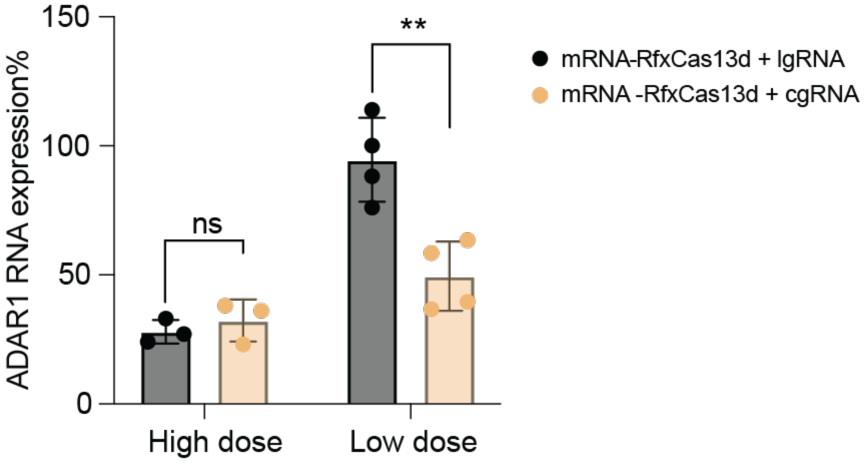


**Figure S3.** *Adar1* knockdown efficiency using mRNA-RfxCas13d (0.5 μg/mL) together with lgRNA or cgRNA (High dose: 0.5 μg/mL; Low dose: 0.25 μg/mL), respectively, in 4T1 cells at different gRNA concentrations for 24 h. RNA was transfected using Lipofectamine 2000.

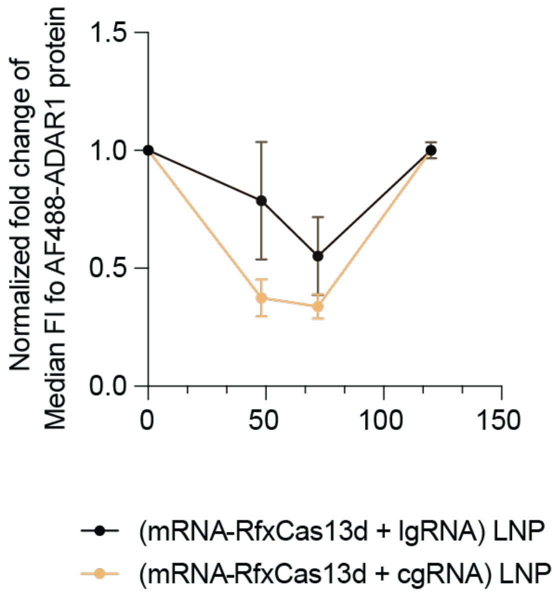


**Figure S4**. ADAR1 protein expression kinetics in MC38 cells before and after LNP co-transfection of mRNA-RfxCas13d together with lgRNA or cgRNA, respectively.

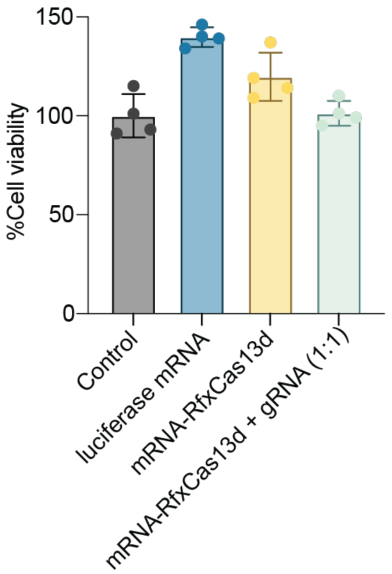


**Figure S5**. 4T1 cell viability 48 h after mRNA-RfxCas13d (0.5 µg/mL) and gRNA (0.5 µg/mL) co-transfection.

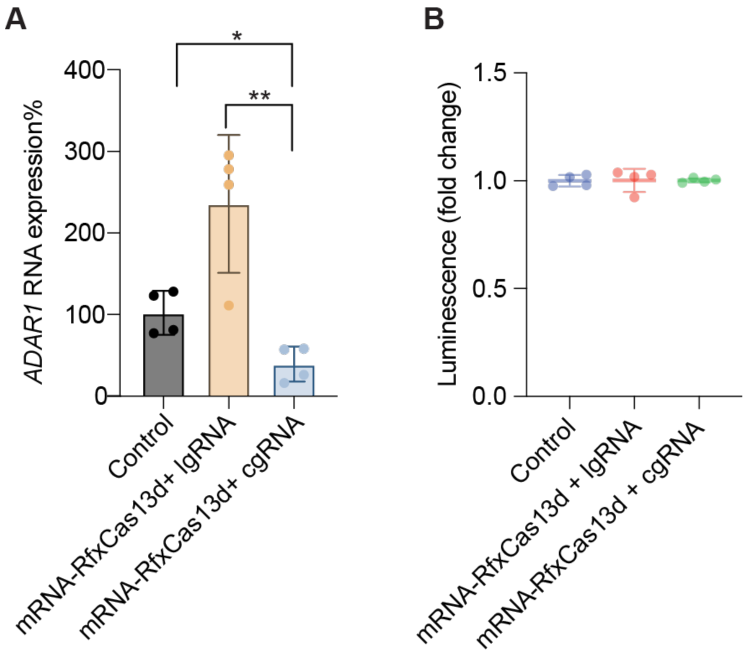


**Figure S6.** mRNA-RfxCas13d + cgRNA showed efficient *ADAR1*-specific knockdown in human cells with undetectable collateral activities. A) *ADAR1* knockdown efficiency in human MDA-MB 231TNBC cells with a luciferase reporter. B) Evaluation of the collateral activity of the above treatments in luciferase-expressing MDA-MB 231TNBC cells via assessing the activities of the luciferase reporter to generate bioluminescence in the presence of D-luciferin substrate. mRNA-RfxCas13d (0.5 µg/mL) and gRNA (0.5 µg/mL) was co-transfected using lipofectamine 2000 for 24 h.

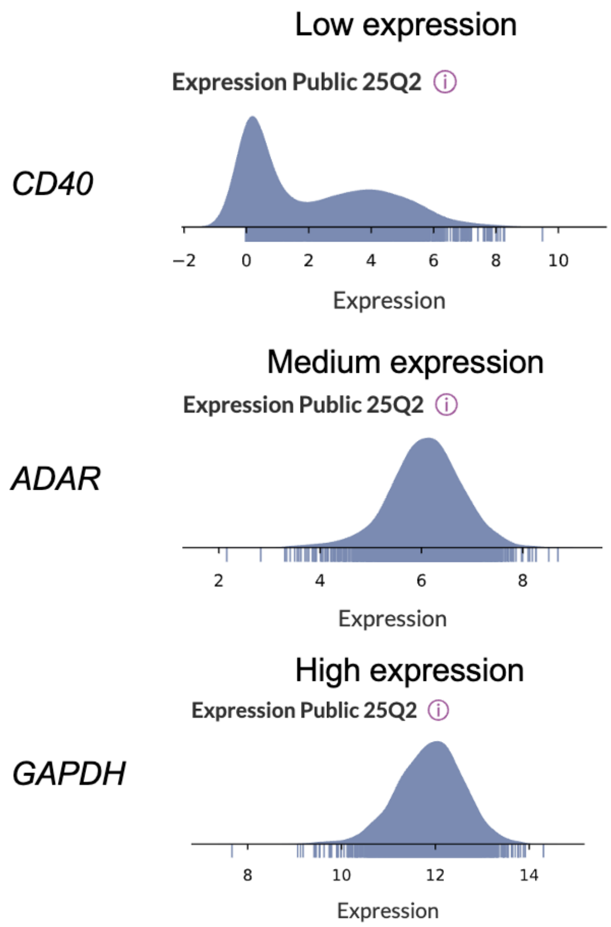


**Figure S7.** Classification of gene expression level in cancer cells from DepMap Public 25Q2 Dataset.

**Table S1.** Characterization of RNA-loaded LNPs by dynamic light scattering. PDI: polydispersity index.

| LNPs | Diameters/nm | PDI |
|---|---|---|
| (mRNA-RfxCas13d + lgRNA) LNPs | 104.1± 18 | 0.14 ± 0.03 |
| (mRNA-RfxCas13d + cgRNA) LNPs | 136.7 ± 29 | 0.18 ± 0.05 |
